# Wild-type FUS corrects ALS-like disease induced by cytoplasmic mutant FUS through autoregulation

**DOI:** 10.1101/2020.12.16.423060

**Authors:** Inmaculada Sanjuan-Ruiz, Noé Govea-Perez, Melissa Mcalonis-Downes, Stéphane Dieterle, Salim Megat, Sylvie Dirrig-Grosch, Gina Picchiarelli, Diana Piol, Qiang Zhu, Brian Myers, Chao-Zong Lee, Don W Cleveland, Clotilde Lagier-Tourenne, Sandrine Da Cruz, Luc Dupuis

## Abstract

Mutations in FUS, an RNA-binding protein involved in multiple steps of RNA metabolism, are associated with the most severe forms of amyotrophic lateral sclerosis (ALS). Accumulation of cytoplasmic FUS is likely to be a major culprit in the toxicity of *FUS* mutations. Thus, preventing cytoplasmic mislocalization of the FUS protein may represent a valuable therapeutic strategy. FUS binds to its own pre-mRNA creating an autoregulatory loop efficiently buffering FUS excess through multiple proposed mechanisms including retention of introns 6 and/or 7. Here, we introduced a wild-type *FUS* gene allele, retaining all intronic sequences, in mice whose heterozygous or homozygous expression of a cytoplasmically retained FUS protein (Fus^ΔNLS^) was previously shown to provoke ALS-like disease or postnatal lethality, respectively. Wild-type FUS completely rescued the early lethality caused by the two *Fus*^ΔNLS^ alleles, and improved age-dependent motor deficit and reduced lifespan associated with the heterozygous expression of *Fus*^ΔNLS^. Mechanistically, wild-type FUS decreased the load of cytoplasmic FUS, increased exon 7 skipping and retention of introns 6 and 7 in the endogenous mouse *Fus* mRNA, leading to decreased expression of the mutant mRNA. Thus, the wild-type *FUS* allele activates the homeostatic autoregulatory loop, maintaining constant FUS levels and decreasing the mutant protein in the cytoplasm. These results provide proof of concept that an autoregulatory competent wild-type FUS expression could protect against this devastating, currently intractable, neurodegenerative disease.

## Introduction

Amyotrophic lateral sclerosis (ALS), the major adult onset motor neuron disease^1,2^, is characterized by a progressive paralysis leading to death within a few years after onset. Mutations in *FUS* cause the most severe cases of ALS, with young onset and rapid disease progression^3,4^. *FUS* mutations are clustered in the C-terminal region of the protein, carrying a PY-nuclear localization sequence (NLS), responsible for its nuclear import. Truncating mutations have been described in ALS families, leading to complete loss of the PY-NLS, and cytoplasmic aggregation of FUS^5,6^. Studies in mouse models have demonstrated that cytoplasmic accumulation of FUS provokes motor neuron degeneration^7–12^. Indeed, heterozygous *Fus* knock-in mice with ALS-like truncating mutations develop mild, late onset muscle weakness and motor neuron degeneration, while haploinsufficient *Fus* knock-out mice do not show ALS related symptoms^10–12^. A successful therapeutic strategy for *FUS*-ALS may lie in reduction of the cytoplasmic FUS content, to avoid its toxic effects.

FUS levels are tightly controlled by autoregulatory mechanisms. Indeed, the addition of more than 20 copies of the complete human *FUS* gene to the mouse genome only slightly increases FUS protein levels, and does not lead to phenotypic consequences^8^, showing the efficacy of this buffering system of FUS levels. Contrastingly, the saturation of FUS autoregulation, through overexpression of cDNA driven, autoregulatory incompetent, FUS expression, is highly toxic to neurons^9,13^. Here, we tested the hypothesis that the expression of a wild-type *FUS* gene, carrying all regulatory elements necessary for autoregulation would engage autoregulation of the mutation carrying RNA, and subsequently decrease accumulation of FUS in the cytoplasm.

## Results

### Wild-type *FUS* transgene rescues lethality and motor defects in *Fus*^ΔNLS^ mice

Human wild-type FUS transgenic mice (hFUS mice) expressing human *FUS* gene including its own human *FUS* promoter obtained from a BAC^8^ were crossed with *Fus*^ΔNLS^ mice^11^ in a two-round mating (**Fig. 1A**). As previously described, *Fus*^ΔNLS/ΔNLS^ mice (in absence of hFUS) die within the first hours after birth^11^ and no mice homozygous mutant *Fus*^ΔNLS^ was obtained at 1 month of age in the absence of hFUS (**Fig. 1B**). Contrastingly, expression of hFUS transgene completely rescued lethality of homozygous *Fus*^ΔNLS/ΔNLS^ mice until adulthood (**Fig. 1C**). About 30% of *Fus*^ΔNLS/+^ mice died before 600 days of age (p=0.0398, log rank, *Fus*^+/+^ vs *Fus*^ΔNLS/+^), similar to what has been observed in mice homozygously expressing another cytoplasmically-retained mutant FUS protein^10^. Most *Fus*^ΔNLS/+^/hFUS mice survived until this age, and their survival rate was indistinguishable from non-transgenic normal mice (p=0.33 *Fus*^+/+^ vs *Fus*^ΔNLS/+^/hFUS) (**Fig. 1D**). The mild, late onset, muscle weakness observed in *Fus*^ΔNLS/+^ mice using inverted grid test^12^, was rescued in *Fus*^ΔNLS/+^/hFUS and in *Fus*^ΔNLS/ΔNLS^/hFUS mice (**Fig. 1E** and **Fig. S1**). Furthermore, hindlimb grip strength deficits associated with expression of *Fus*^ΔNLS/+^ were mildly and transiently improved in *Fus*^ΔNLS/+^/hFUS females (**Fig. 1F**) but not in males (**Fig. 1F**). Indeed, in this test, the performance of hFUS transgenic mice decreased significantly in males after 10 months of age, thus confounding a potential protection (**Fig. 1F** and **Fig. S1**). Thus, wild-type human FUS significantly rescued lethality and, at least partially, motor deficits associated with cytoplasmically retained mutant *Fus*^ΔNLS^ protein.

**Figure 1:**
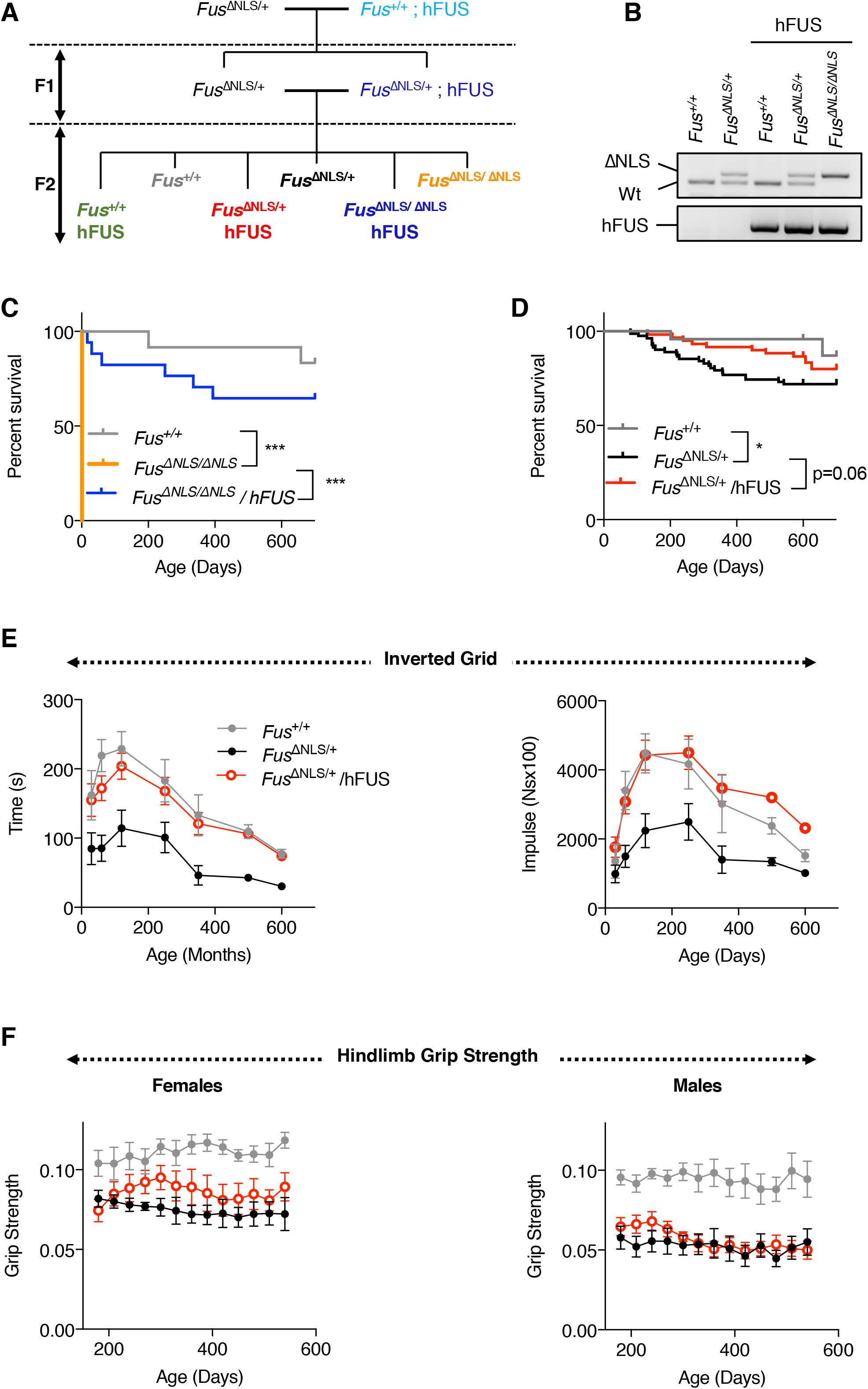
hFUS transgene rescues lethality and motor deficits in *Fus*^ΔNLS^ mice. **A:** Scheme of the breeding strategy **B**: Representative genotyping results of 5 mice at 1 month of age. **C-D**: Kaplan Meier survival curve of the different genotypes either heterozygous (C) or homozygous for the ΔNLS mutation (D). Note that all *Fus*^ΔNLS/ΔNLS^ mice die at birth, unless carrying a hFUS transgene. *, p<0.05 Log Rank test; ***, p<0.001 Log rank test **E**: Age-dependent changes in the mean hanging time (s) and holding impulse (N s) in the four-limb wire inverted grid test in *Fus*^+/+^, and *Fus*^Δ*NLS*/+^ mice with or without hFUS transgene. *N* =10-28 per group. Mixed effect analysis, with 3 factors (Age, ΔNLS genotype and hFUS genotype). *P<0.001* for ΔNLS genotype, *P<0.001* for age, *P<0.001* for hFUS genotype. A significant protective interaction is observed between ΔNLS and hFUS genotypes (p=0.0216, and *p=0.0366).* Only 3 groups out of 5 are shown here for clarity. The whole dataset is shown in Fig S1. **F:** Hindlimb grip strength in female and male mice. Mixed effect analysis, with 3 factors (Age, ΔNLS genotype and hFUS genotype). For female mice, P<0.001 for ΔNLS genotype, p=ns for age, p=ns for hFUS genotype. A significant protective interaction is observed between ΔNLS and hFUS genotypes (p=0.0131). For male mice (H), P<0.001 for ΔNLS genotype, p=ns for age, p=ns for hFUS genotype. No significant protective interaction is observed between ΔNLS and hFUS genotypes (p=0.0512). Only 3 groups out of 4 are shown here for clarity. The whole dataset is shown in Fig. S1.

### Wild-type *FUS* transgene decreases cytoplasmic accumulation of FUS in *Fus*^ΔNLS^ mice

We then asked whether hFUS transgene altered subcellular localization of FUS in *Fus*^ΔNLS^ mice. As expected^11,12^, cytoplasmic FUS levels were elevated by five-fold in cerebral cortex of *Fus*^ΔNLS/+^ mice as compared to corresponding wild-type mice (**Fig. 2A** and **Fig. S2-3** for quantification and uncropped western blots). This was not observed when an antibody targeting the NLS sequence (C-term FUS), absent from the Fus^ΔNLS^ protein, was used, demonstrating that this increase is related to the mislocalization of the mutant protein. Importantly, the increase in mouse FUS in cytoplasmic fractions of *Fus*^ΔNLS/+^ mice, was normalized by hFUS transgene (**Fig. 2A**). Contrastingly, nuclear FUS levels were similar in all genotypes, irrespective of the presence of the *Fus*^ΔNLS^ mutation or that of the hFUS transgene. Human FUS levels were increased in *Fus*^ΔNLS/ΔNLS^ mice carrying a hFUS transgene, likely compensating for the loss of nuclear FUS of mouse origin. In spinal cord sections, *Fus*^ΔNLS/+^ neurons displayed a mixed cytoplasmic and nuclear FUS staining, that was prevented by the hFUS transgene (**Fig. 2B**), and this was also observed in motor neurons as identified using double FUS/ChAT immunofluorescence (**Fig. 2C**). Accumulation of cytoplasmic asymmetrically demethylated (ADMA) FUS is a feature of *FUS*-ALS^5,6,14^ patients which was recapitulated in the *Fus*^ΔNLS/+^ mice, as we previously reported^12^. Here, this significant increase in ADMA-FUS detected in *Fus*^ΔNLS/+^ cytoplasmic fractions, was largely prevented by the hFUS transgene in *Fus*^ΔNLS/+^/hFUS mice (**Fig. 2A**), but not in *Fus*^ΔNLS/ΔNLS^/hFUS mice. While ADMA-FUS immunoreactivity was diffuse in the cytoplasm of *Fus*^ΔNLS/+^ motor neurons, expression of the hFUS transgene in *Fus*^ΔNLS/+^/hFUS and *Fus*^ΔNLS/ΔNLS^/hFUS mice led to more localized, perinuclear ADMA-FUS immunoreactivity in *Fus*^ΔNLS/+^ mice (**Fig. 2D**), thus suggesting that wild-type hFUS restores aberrant FUS nearly to normal levels.

**Figure 2:**
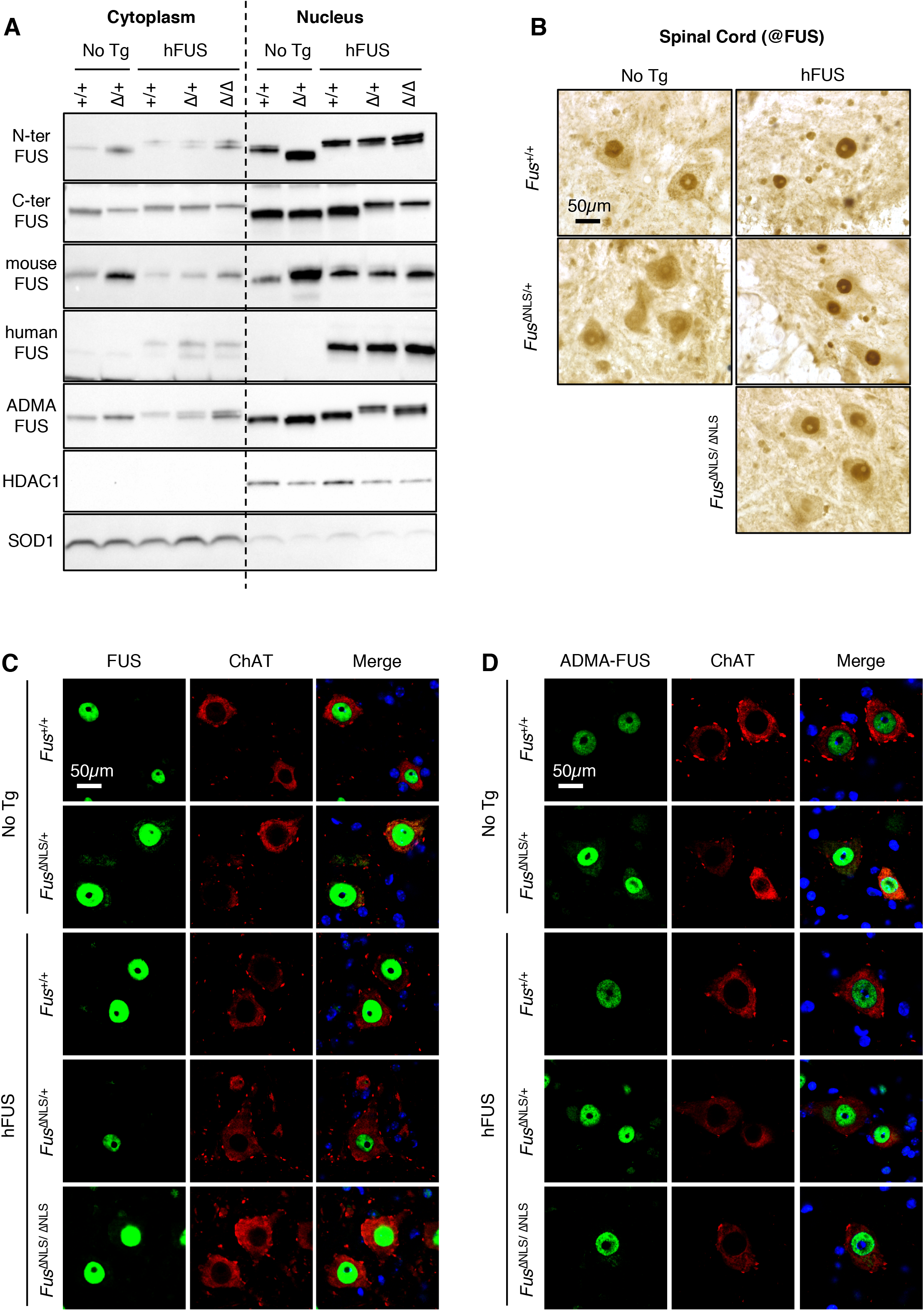
hFUS transgene corrects abnormal cytoplasmic accumulation of FUS protein in *Fus*^ΔNLS^ mice. **A**: Immunoblot analysis of FUS protein subcellular localization in cortex of *Fus*^+/+^ (+/+) and *Fus*^*ΔNLS*/+^ (Δ/+) mice with or without hFUS transgene and of *Fus*^*ΔNLS*/ΔNLS^ mice with hFUS transgene at 1 month of age. Representative results using different antibodies targeting the N-terminal part (N-ter) of FUS, the C-terminal (C-ter) NLS, mouse FUS, human FUS and asymmetrically arginine dimethylated FUS (ADMA-FUS). SOD1 and HDAC1 are used as loading controls for cytoplasmic and nuclear protein extracts fractions, respectively. Note that these immunoblots were performed on different membranes to avoid cross reaction between different antibodies. Uncropped western blots and corresponding stain free gels are provided in Figure S3. **B**: FUS immunohistochemistry in the spinal cord ventral horn at 22 months of age of mice of the indicated genotypes. Note that diffuse FUS cytoplasmic staining, obvious in *Fus*^*ΔNLS*/+^ mice, is rescued by the hFUS transgene. Scale bar: 50μm **C-D:** Double immunostaining for the motoneuronal marker ChAT and N-terminal FUS (D), or ChAT and ADMA-FUS in the spinal cord ventral horn at 22 months of age. Note the cytoplasmic redistribution of truncated FUS in *Fus*^*ΔNLS*/+^ mice and its rescue by hFUS transgene. Scale bar: 50μm

### Wild-type *FUS* transgene activates autoregulation of mutant *Fus* to decrease mutant FUS protein

The increase in *Fus* mRNA in *Fus*^ΔNLS/+^ mice was fully corrected by expression of wild-type hFUS transgene in 1-month old spinal cord (**Fig. 3A-B**) and frontal cortex (**Fig. S4**) of *Fus*^ΔNLS/+^/hFUS and *Fus*^ΔNLS/ΔNLS^/hFUS animals, leading to accumulated mouse *Fus* mRNA levels close to those of endogenous *Fus* in normal non-transgenic mice. This restoration of mouse *Fus* mRNA levels by hFUS transgene was sustained through aging as observed in 22-month old *Fus*^ΔNLS/+^/hFUS mice. Consistently, mutant *Fus*^Δ*NLS*^ mRNA levels decreased in spinal cord and frontal cortex of *Fus*^ΔNLS/+^/hFUS and *Fus*^ΔNLS/ΔNLS^/hFUS animals compared to the *Fus*^ΔNLS/+^ mice (**Fig. 3D** and **Fig. S4**), while human *FUS* mRNA levels remained comparable across the three genotypes with hFUS transgene (**Fig. 3C** and **Fig. S4**).

**Figure 3:**
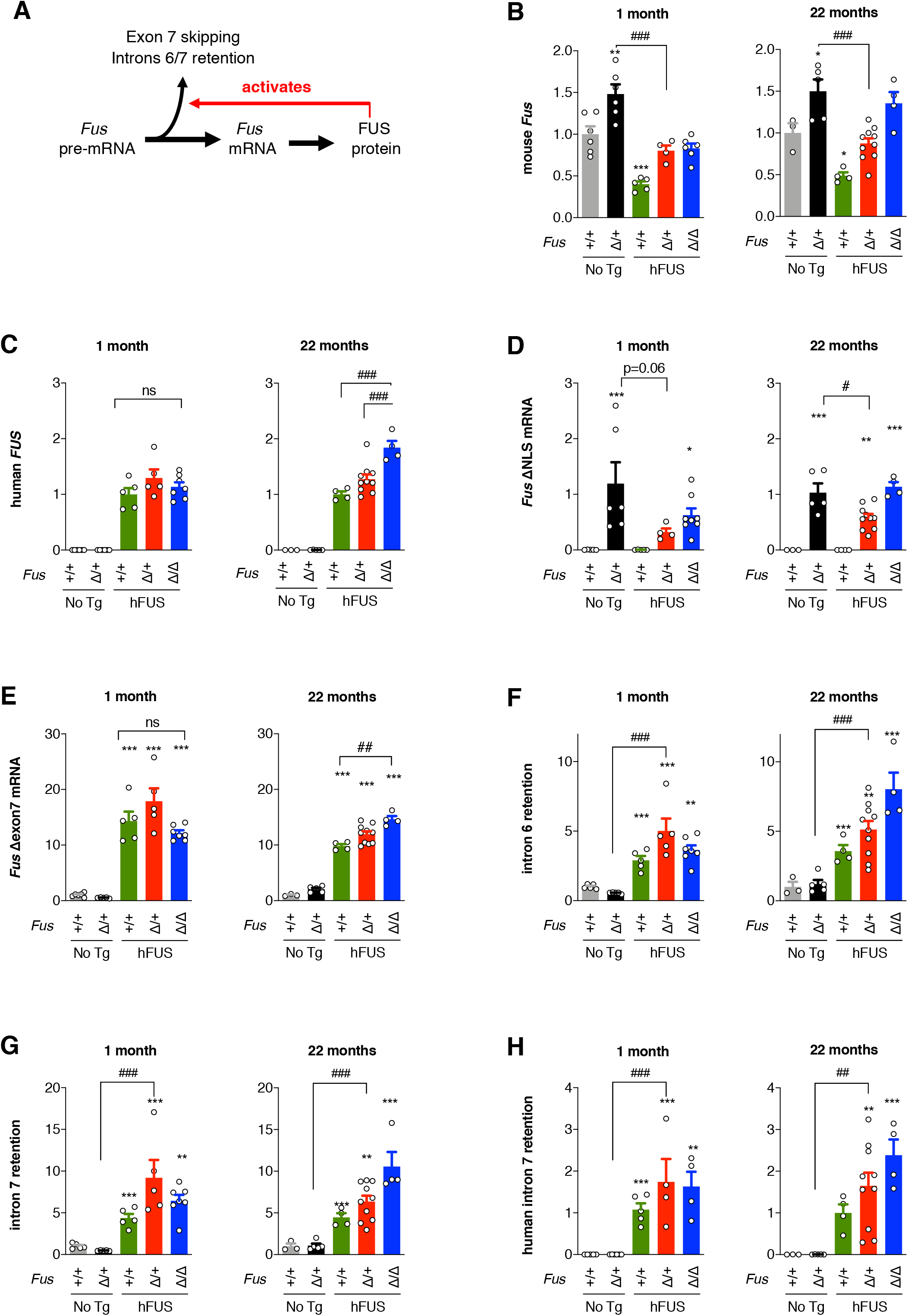
hFUS transgene downregulates endogenous Fus mRNA levels and activates autoregulatory splicing in *Fus*^ΔNLS/+^ spinal cord. **A:** Scheme depicting autoregulatory pathway of FUS expression **B-H**: RT-qPCR results for endogenous mouse *Fus* mRNA (B), human FUS transgene (C) and mutant *Fus* mRNA carrying the ΔNLS mutation in spinal cord at 1 month of age (D), endogenous *Fus* mRNA deleted of exon 7 (E), endogenous *Fus* mRNA retaining intron 6 (F), endogenous *Fus* mRNA retaining intron 7 (G) and exogenous *FUS* mRNA retaining intron 7 (H) in spinal cord at 1 month of age (left) or 22 months of age (right). Note that the hFUS transgene decreases expression of endogenous *Fus* gene and leads to decreased expression of mutant *Fus* mRNA at 1 and 22 months of age. Furthermore, the hFUS transgene activates autoregulatory exon 7 skipping as well as retentions of introns 6 and 7 in endogenous mRNA and retention of intron 7 in exogenous mRNA at 1 and 22 months of age. *N* = 4-8. *, p<0.05, \*\**p* < 0.01, \*\**p* < 0.001 vs *Fus*^*+/+*^, #, p<0.05, ###, p<0.001 vs indicated genotype by ANOVA followed by Tukey.

We further investigated the three possible autoregulatory mechanisms that have been documented for FUS. First, FUS protein is proposed to bind to its own pre-mRNA, leading to the splicing of exon 7, and the possible subsequent degradation of the abnormally Δexon 7 *FUS* mRNA through nonsense-mediated mRNA decay^15,16^. Interestingly, expression of hFUS transgene increased levels of the aberrantly spliced *Fus* Δexon 7 mRNA (**Fig. 3E** and **Fig. S4**). Secondly, increased FUS levels have recently been reported to lead to the retention of introns 6 and 7 in the mature mRNA, and to the nuclear retention of the aberrant transcripts^17^. *Fus* endogenous mRNAs with retained introns 6 or 7 strongly increased in all mice expressing hFUS transgene at 1- and 22-months of age (**Fig. 3F-G** and **Fig. S4**). We also observed prominent retention of human intron 7 in all samples derived from mice expressing the hFUS transgene (**Fig. 3H** and **Fig S4**). Thirdly, besides intron skipping and retention, FUS was also found to regulate its own levels through the stimulation of miR200^18^. Another target of miR200 is ZEB1, whose expression is dependent upon levels of miR200^19,20^. *Zeb1* expression appears unchanged in *Fus*^ΔNLS/+^ tissues, whether or not expressing the h*FUS* transgene (**Fig. S5**), indirectly suggesting that this latter autoregulatory mechanism is not engaged in the effects mediated by the hFUS transgene. Thus, wild-type human FUS gene decreases expression of the endogenous *Fus* gene through increased retention of introns 6 and 7 and skipping of exon 7, leading to decreased production of toxic *Fus*^ΔNLS^ protein, and subsequent alleviation of all the downstream consequences of the expression of cytoplasmically mislocalized mutant FUS.

## Discussion

In the current study, we show that providing a wild-type allele of the *FUS* gene is sufficient to rescue ALS-like phenotypes associated with cytoplasmically retained mutant FUS protein expression.

Our result appears *a priori* paradoxical since the toxicity of FUS mutations was shown to be largely driven by cytoplasmic Fus^7–12^, that is not expected to be directly compensated by the wild-type protein. Furthermore, overexpression of the wild-type protein was shown to be toxic to neurons^9,13,21^. Normal nuclear FUS levels in *Fus*^Δ*NLS/+*^ mice indicate that the hFUS transgene is not simply rescuing loss of nuclear FUS. On the contrary, the hFUS transgene appears to indirectly protect from accumulation of mutant protein through the autoregulatory loop maintaining nuclear FUS levels^15–17^. Indeed, nuclear FUS levels are stable in all groups, suggesting homeostatic mechanisms to avoid the toxicity of loss of nuclear FUS^22–26^ or its excess ^9,13,21^. FUS autoregulation is positively demonstrated by variations in *Fus* mRNA levels in our different experimental conditions. First, *Fus* mRNA levels are increased in *Fus*^Δ*NLS/+*^ mice, thereby compensating the proportion of FUS protein translated from the mutant allele and unable to enter the nucleus. Conversely, in human *FUS* transgenic mice the addition of the exogenous *FUS* transgene is sufficient to decrease endogenous mouse *Fus* mRNA levels, consistent with previous studies^8,9^. In our study, the addition of the hFUS transgene in *Fus*^Δ*NLS/+*^ mice rescued *Fus* overexpression in *Fus*^Δ*NLS/+*^ mice, and decreased mutant mRNA levels. Since this overexpression acts as a feed forward mechanism amplifying the cytoplasmic accumulation of FUS, avoiding this overexpression might on its own be sufficient to slow down the vicious cycle leading to phenotypes in *Fus*^Δ*NLS/+*^ mice. Importantly, the exogenous human transgene is also, on its own, subject to autoregulation, and this likely explains why genomic based constructs are much less toxic than cDNA-based constructs devoid of required autoregulatory sequences.

Our results suggest that gene therapy to reintroduce the wild-type protein, while including sequences required for autoregulation, would enable the correction of molecular and behavioral phenotypes, meanwhile avoiding the toxicity of wild-type protein overexpression in *FUS-*ALS. Besides *FUS*-ALS, FUS mutations have been associated with other neurodegenerative diseases, such as frontotemporal dementia^27,28,29^, chorea^30^, mental retardation^31^, psychosis^32^ and essential tremor^33^. FUS aggregation has been observed in sporadic ALS^34–36^ and FTD^37–39^, but also in spino-cerebellar ataxia and Huntington’s disease^40,41^. A gene therapy to restore normal nuclear FUS levels might thus be relevant for other patients to be identified. Last, it is noteworthy that similar autoregulatory mechanisms exist for other RNA-binding proteins, in particular TDP-43^42–47^ or hnRNPA1^48^. Whether utilizing such autoregulatory mechanisms to decrease mutant protein through overexpression of a wild-type protein might be a general therapeutic approach in such diseases remains to be determined.

## Materials and methods

### Mouse models and genotyping

Mouse experiments were approved by local ethical committee from Strasbourg University (CREMEAS) under reference number 2016111716439395 and all experimental procedures performed in San Diego were approved by the Institutional Animal Care and Use Committee of the University of California, San Diego. Transgenic mice were generated as described in ^11,12^ and ^8^, were bred in Charles River animal facility and housed in the Faculty of medicine from Strasbourg University with 12/12 hours of light/dark cycle (light on at 7:00 am) under constant conditions (21 ± 1 °C; 60% humidity) and with unrestricted access to food and water.

Mice were weaned and genotyped at 21 days by PCR from tail biopsy, or at death if occurring before 21 days of age.

The following primer sequences were used to genotype mice:

hFUS-For: GAATTCGTGGACCAGGAAGGTC
hFUS-Rev: CACGTGTGAACTCACCGGAGTCA
FUS-For: GAT-TTG-AAG-TGG-GTA-GAT-AGT-GCA-GG
FUS-Rev: CCT-TTC-CAC-ACT-TTA-GTT-TAG-TCA-CAG

Heterozygous Fus knock-in mice, lacking the PY-NLS, were crossed with mice expressing human wild type FUS from a complete, autoregulatory competent, human gene to obtain following genotypes: *Fus*^+/+^, *Fus*^Δ*NLS*/+^, *Fus*^Δ*NLS*/ΔNLS^, *Fus*^+/+^/hFUS, *Fus*^ΔNLS/+^/hFUS, *Fus*^ΔNLS/ΔNLS^/hFUS. The genetic background of all mice used in this study is C57Bl6/J. Breeding steps were performed in parallel in both laboratories. 76 mice of the F2 generation were generated in Strasbourg, and 110 mice of the F2 generation were generated in San Diego.

### Mouse behavior

#### Survival

Survival was studied during the first hours after birth and dead new born mice were genotyped. Mice surviving the post-natal period were genotyped at 21 days and followed weekly until death or euthanized using ketamine-xylazine when they reach the following endpoints: auto-mutilation, weight loss greater than 10% of the initial weight and when they could not turn around again within 10 seconds after being laid on their side.

#### Inverted grid

Mice were habituated for 30 min in the test room prior testing. Motor performance were performed weekly as described previously ^12^ from 1 month until 22 months of age. The wire grid hanging time (or “hang time”) was defined as the amount of time that it takes the mouse to fall down from the inverted grid and was measured visually with a stopwatch. The procedure was repeated 3 times during 5min with 5 min break between tests. All mice were returned to their homecage after completing the test. The holding impulse corresponds to hanging time normalized with mouse weight and gravitational force.

#### Grip test

Grip strength was measured using a Grip Strength Meter (Columbus Instruments, Columbus, OH) on cohorts (N=12-30) made up of approximately the same number of males and females. Mice were allowed to grip a triangular bar only with hind limbs, followed by pulling the mice until they released; five force measurements were recorded in each separate trial.

### Histological techniques

Mice aged of 1 month or 22 months were anesthetized with intraperitoneal injection of 100 mg/kg ketamine chlorhydrate and 5mg/kg xylazine then perfused with PFA 4%. After dissection, spinal cord was included in agar 4% and serial cuts of 40μm thick were made with vibratome.

#### Peroxidase immunohistochemistry

For peroxidase immunohistochemistry, sections were incubated 10 min with H_2_O_2_ 3%, washed 3 times and blocked with 8% Horse serum, 0,3% Bovine Serum Albumin and 0,3% Triton in PBS with 0,02% Thimerosal. Sections were incubated with rabbit anti-FUS antibody (ProteinTech 11570-1-AP; diluted 1:100) in blocking solution overnight at room temperature. After washing sections, they were incubated for 2h at room temperature with biotinylated donkey anti-rabbit antibody (Jackson 711-067-003; diluted 1:500) in blocking solution. Then, sections were washed, incubated for 1h in horseradish peroxidase ABC kit (Vectastain ABC kit, PK-6100, Vector Laboratories Inc.), washed and incubated with DAB (Sigma, D5905). The enzymatic reaction was stop by adding PBS 1X and washed with water. Finally, sections were mounted with DPX mounting medium (Sigma, O6522).

#### Immunofluorescence

Sections were incubated in blocking solution (8% Horse serum, 0.3% Bovine Serum Albumin, 0.3% Triton, PBS-0.02% Thimerosal) at room temperature for 1 hour, then incubated overnight at room temperature in primary antibody with blocking solution: rabbit anti-FUS antibody (ProteinTech, 11570-1-AP, 1:100), goat anti-ChAT (Millipore, AB144P, 1/50), rat anti-ADMA (^5,6^, kind gift of Pr C. Haass, Munich Germany, 1/100) or rabbit anti-Ubi (Abcam, ab179434, 1/100). After 3 rinses in PBS, sections were incubated for 2h at room temperature with Hoechst (Sigma, B2261, 1/50.000) and secondary antibodies in blocking solution: Alexa-488-conjugated donkey anti-rabbit secondary antibody (Jackson, 711-547-003, 1/500) or Alexa-594-conjugated donkey anti-goat secondary antibody (Molecular Probes, A 11058, 1/500). Finally, sections were subsequently washed with PBS 1X (3 × 10 min) and mounted in Aqua/polymount (Polysciences 18606).

### Tissue fractionation and western blotting

Tissues were washed in PBS1x and lysed in NE-PER Nuclear and Cytoplasmic Extraction (Thermo Scientific, 78835) according to the manufacturer’s instructions. Protein extracts were dosed by BCA Assay (Interchim, UP95424A, UP95425A). Thereafter proteins were denatured and SDS page was performed with 30μg of cytoplasmic proteins and 10μg of nuclear proteins on criterion TGX stain free gel 4-20% (Biorad, 5678094). Proteins were blotted on nitrocellulose membrane using semi-dry Transblot Turbo system (BioRad, France) and blocked with 10% non-fat milk during 1h. Primary antibodies (Rabbit anti-hFUS (^8,9^, #14080, 1/2000), Rabbit anti-mFUS (^8,9^, #14082, 1/4000), Rat anti-FUS ADMA (^5,6^, kind gift of Pr C. Haass, Munich Germany, 1/500), Rabbit anti-FUS 293 (Bethyl, A-300-293A, 1/2000), Rabbit anti-FUS 294 (Bethyl, A300-294A, 1/2000), Sheep anti-SOD1 (Calbiochem, 574597, 1/1000), Rabbit anti-HDAC1 (Bethyl, A300-713A, 1/1000)) were incubated overnight at 4°C in 3% non-fat milk. Washing was proceeded with washing buffer (Tris pH 7.4 1 M, NaCl 5M, Tween 20 100 %) and secondary antibodies (anti-rabbit HRP (PARIS, BI2407,1/5000), anti-sheep HRP (Jackson, 713-035-147, 1/5000)) were incubated 1h30 at room temperature. After successive washes, proteins were visualized with chemiluminescence using ECL Lumina Forte (Millipore, France) and chemiluminescence detector (Bio-Rad, France). Total proteins were detected with stain free gel capacity (Biorad, 5678094) and used to normalize for protein loading. All values were normalized against nuclear levels of FUS in *Fus*^+/+^ extracts set to 1.

### RNA extraction and RT-qPCR

Total RNA was extracted from spinal cord and frontal cortex using TRIzol® reagent (Life Technologies). 1 μg of RNA was reverse transcribed with iScript™ reverse transcription (Biorad, 1708841). Quantitative polymerase chain reaction was performed using Sso Advanced Universal SYBR Green Supermix (Bio-Rad 1725274) and quantified with Bio-Rad software. Gene expression was normalized by calculating a normalization factor using actin, TBP and pol2 genes according to GeNorm software (Vandesompe et al 2002). Primer sequences are provided in **Table S1.**

### Statistics

All results from analysis are presented as mean ± standard error of the mean (SEM) and differences were considered significant when p < 0.05. Significance is presented as follows: * p<0.05, ** p<0.01, and *** p<0.001. For comparison of two groups, two-tailed unpaired Student’s t –test was used in combination with F-test to confirm that the variances between groups were not significantly different. For longitudinal analysis of behavioral data, results were analyzed using a mixed effect analysis with three factors (ΔNLS genotype, hFUS genotype and age) as indicated in the figure legends. Data were analyzed by using the GrapPad Prism version 8.0.

## Acknowledgements

We thank Pr Christian HAASS for the kind gift of the anti-ADMA FUS antibody. This work was funded by Agence Nationale de la Recherche (ANR-16-CE92-0031 to LD, ANR-16-CE16-0015, ANR-19-CE17-0016), by Fondation pour la recherche médicale (FRM, DEQ20180339179 and post-doctoral position to SM), Axa Research Funds (rare diseases award 2019, to LD), Fondation Thierry Latran (HypmotALS, to LD), MNDA (Dupuis/Apr16/852-791 to LD), ALSA (2235, 3209 and 8075 to LD and CLT), Association Francaise de Recherche sur la sclérose latérale amyotrophique (2016, to LD), Target ALS (to CLT) and Muscular Dystrophy Association (to SDC). LD is USIAS fellow 2019. CLT is the recipient of the Araminta Broch-Healey Endowed Chair in ALS. ISR was funded by the Région Grand Est (France).

---

**Supplementary figure 1: hFUS transgene rescues homozygous Fus^ΔNLS/ΔNLS^ mice**

**A:** Age-dependent changes in the mean hanging time (s) and holding impulse (N s) in the four-limb wire inverted grid test in *Fus*^+/+^, and *Fus*^*ΔNLS*/ΔNLS^ mice with hFUS transgene.

**B:** Hindlimb grip strength in female and male mice.

*N* =10-28 per group.

Experimental data for the control group are identical to Figure 2, as mice were littermates and followed simultaneously. Other genotypes were omitted from the figure for clarity.

**Supplementary figure 2: quantification of western blotting experiments of Figure 2A.** Quantification of Nter, C-ter, mouse, human and ADMA-FUS protein levels in cytoplasmic and nuclear fractions of the indicated genotypes. *N* = 4-8. \**p* < 0.05, \*\*\**p* < 0.001vs *Fus*^*+/+*^, #, p<0.05 and ###, p<0.001 vs indicated genotype by ANOVA followed by Tukey.

**Supplementary Figure 3: raw western blotting results.**

Uncropped western blot of Figure 2A, along with the respective StainFree images showing equal loading and used to normalize quantifications.

**Supplementary Figure 4: hFUS transgene downregulates endogenous *Fus* mRNA levels in Fus^ΔNLS/+^ frontal cortex**

**A:** Scheme depicting autoregulatory pathway of FUS expression

**B-H**: RT-qPCR results for endogenous mouse *Fus* mRNA (B), human FUS transgene (C) and mutant *Fus* mRNA carrying the ΔNLS mutation in spinal cord at 1 month of age (D), endogenous *Fus* mRNA deleted of exon 7 (E), endogenous *Fus* mRNA retaining intron 6 (F), endogenous *Fus* mRNA retaining intron 7 (G) and exogenous *FUS* mRNA retaining intron 7 (H) in frontal cortex at 1 month of age (left) or 22 months of age (right).

Note that the hFUS transgene decreases expression of endogenous *Fus* gene and leads to decreased expression of mutant *Fus* mRNA at 1 and 22 months of age. Furthermore, the hFUS transgene activates autoregulatory exon 7 skipping as well as retentions of introns 6 and 7 in endogenous mRNA and retention of intron 7 in exogenous mRNA at 1 and 22 months of age.

*N* = 4-8. *, p<0.05, \*\**p* < 0.01, \*\**p* < 0.001 vs *Fus^+/+^*, #, p<0.05, ###, p<0.001 vs indicated genotype by ANOVA followed by Tukey

**Supplementary Figure 5: *Zeb1* expression in *Fus*^ΔNLS/+^ spinal cord and frontal cortex**

RT-qPCR results for *Zeb1* mRNA in frontal cortex (upper row) and spinal cord (lower row) at 1 month of age (left) or 22 months of age (right).

*N* = 4-8. No significant change in mRNA levels is observed. ANOVA followed by Tukey.

