## Supplementary Figures S1-S5 for "Wild-type FUS corrects ALS-like disease induced by cytoplasmic mutant FUS through autoregulation"

### Sanjuan-Ruiz et al. Figure S1

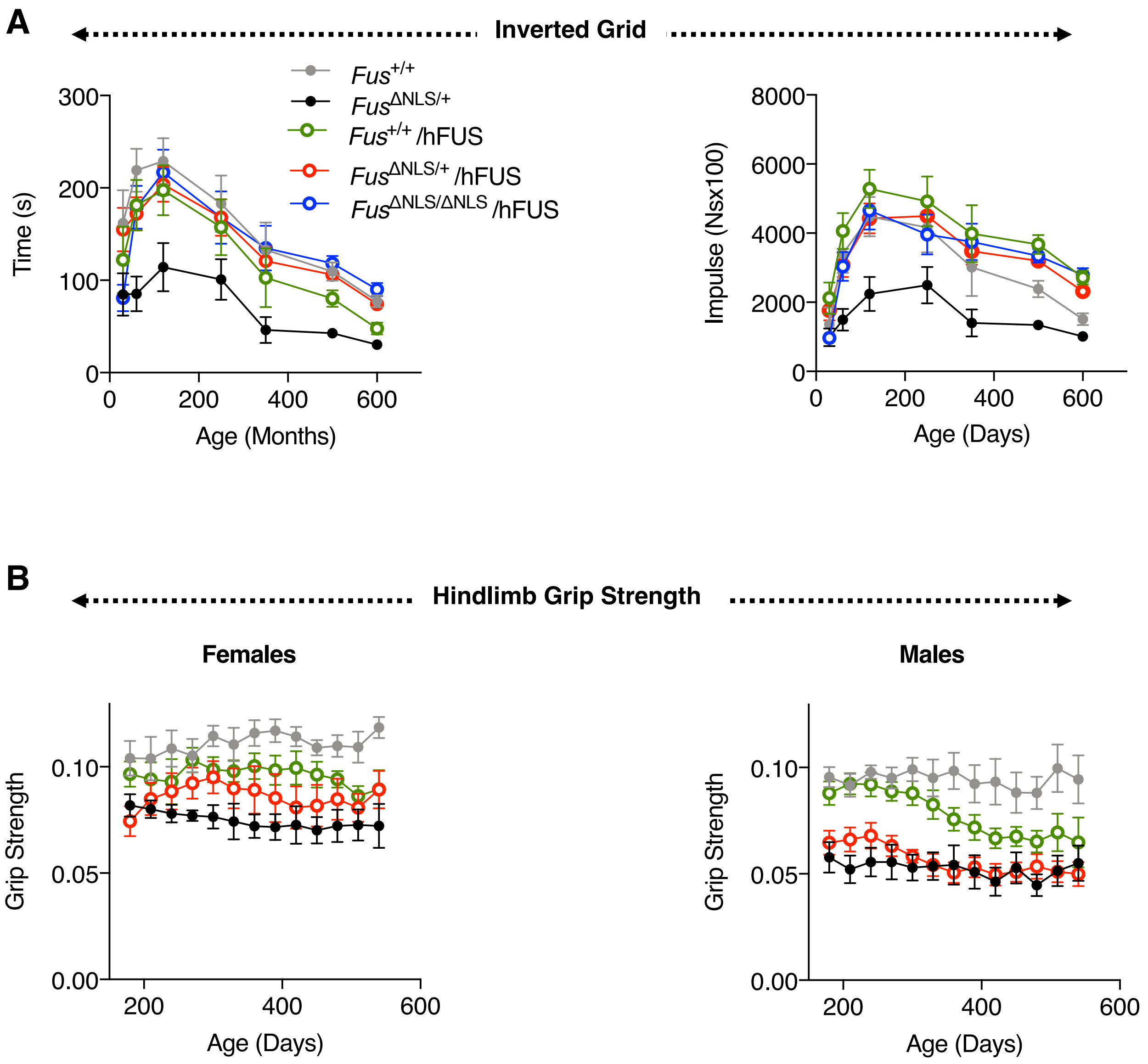

### Sanjuan-Ruiz et al. Figure S2

#### Frontal cortex, 1mo

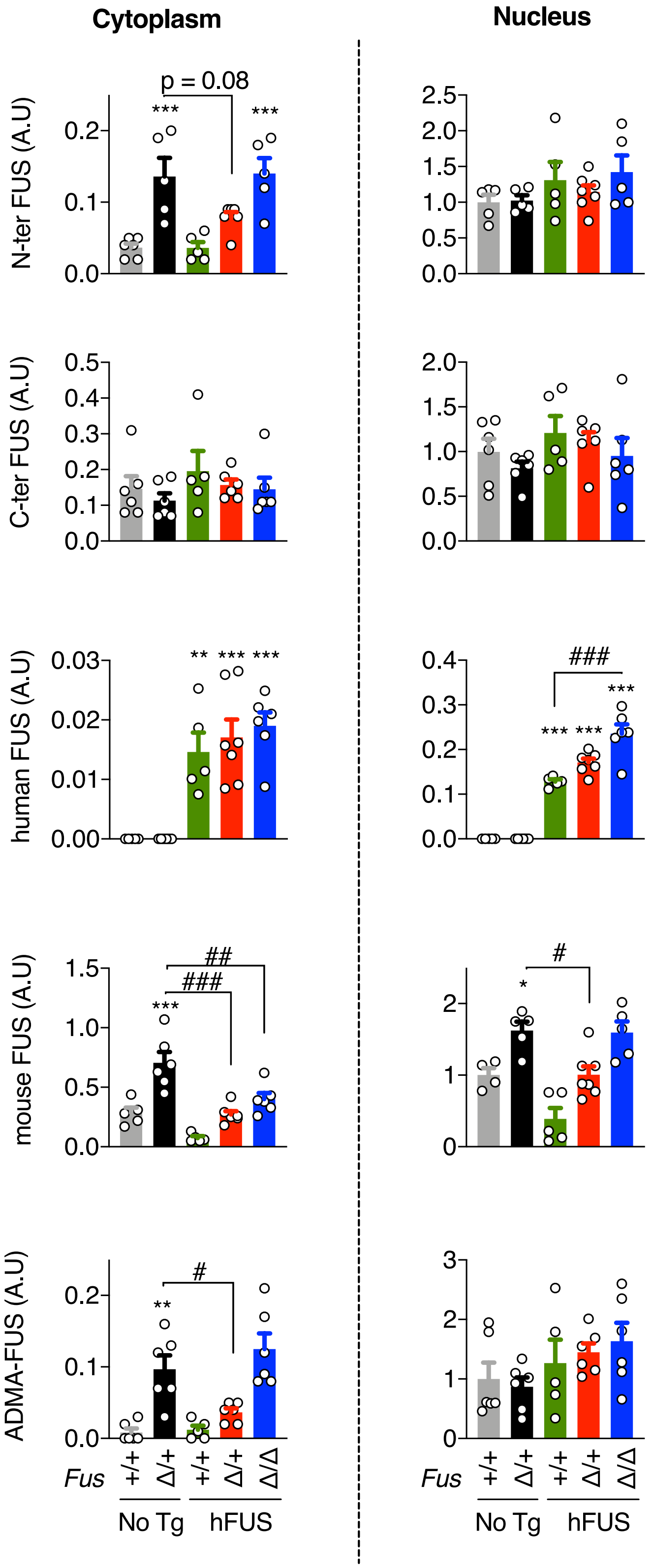

### Sanjuan-Ruiz et al. Figure S3

**Uncropped western blots Figure 2A**

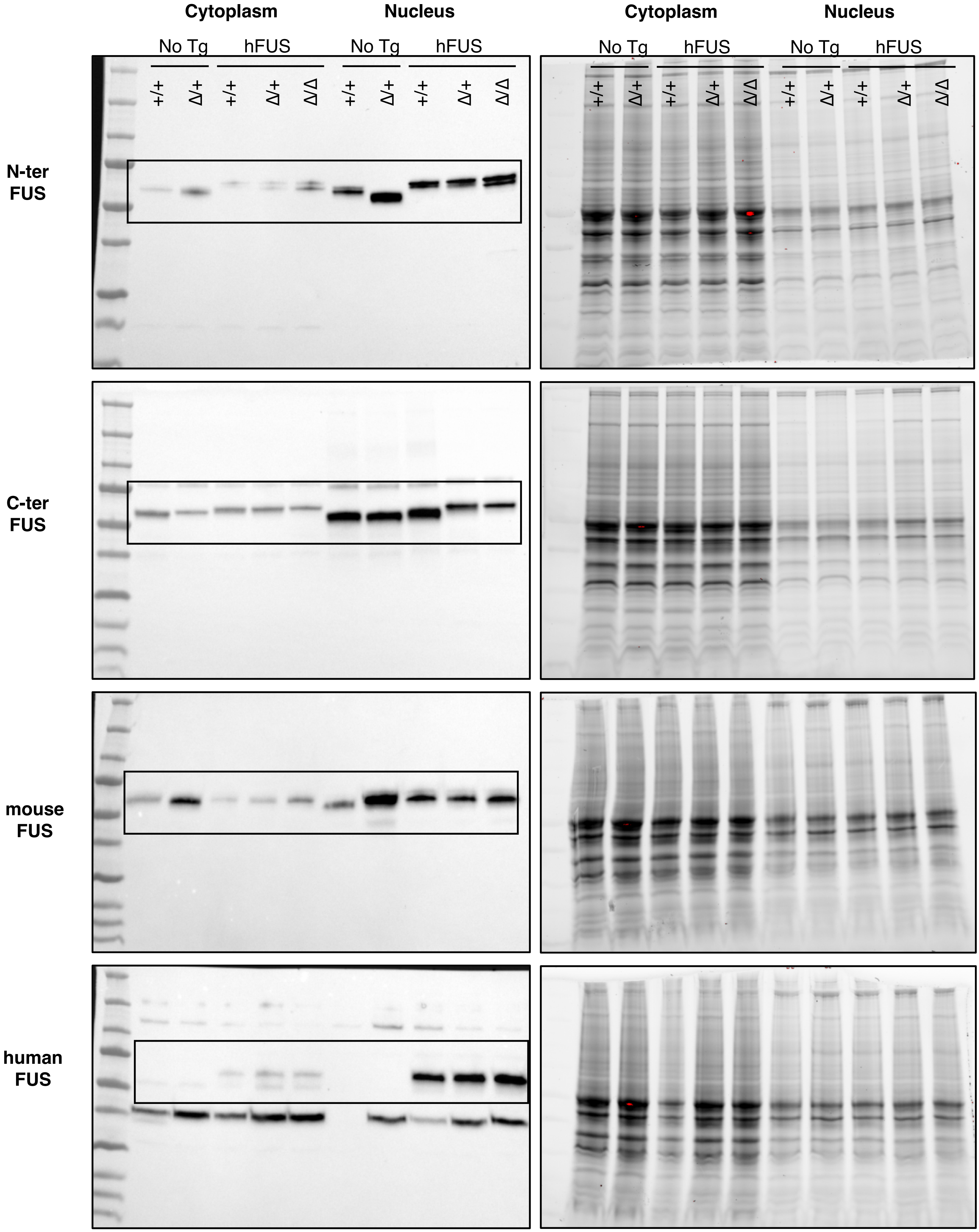

### Sanjuan-Ruiz et al. Figure S3 (continued)

**Uncropped western blots Figure 3A**

**Cytoplasm** **Nucleus**

No Tg hFUS No Tg hFUS

**HDAC1**

**SOD1**

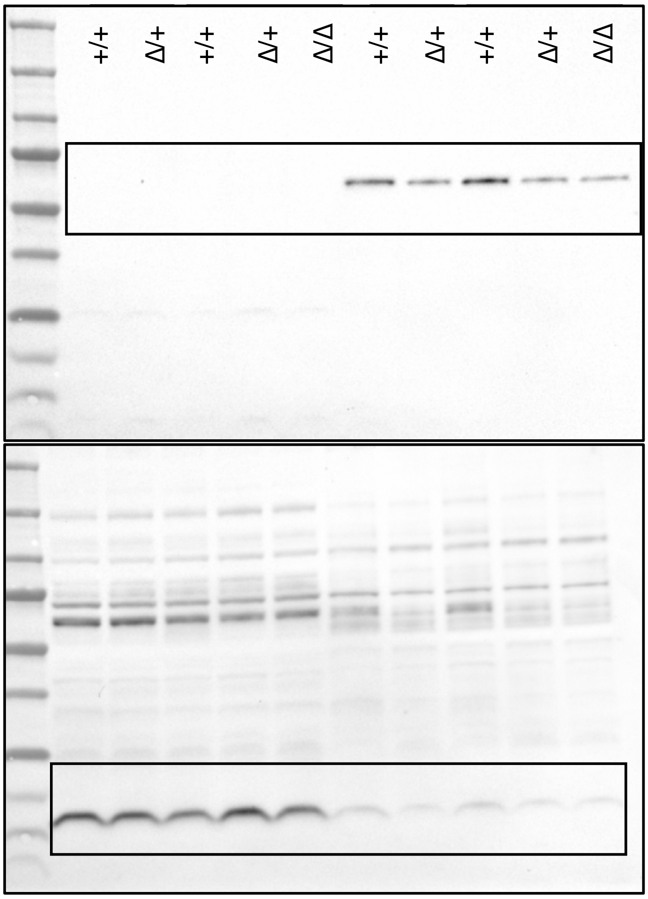


**Cytoplasm** **Nucleus**

No Tg hFUS No Tg hFUS

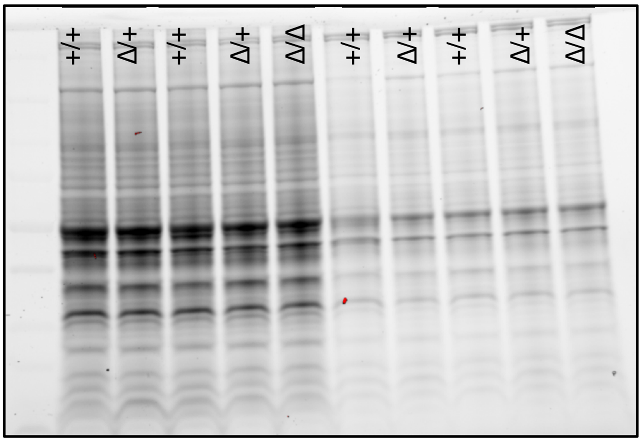


**Cytoplasm** **Nucleus**

No Tg hFUS No Tg hFUS

+/+ Δ/+ +/+ Δ/+ Δ/Δ +/+ Δ/+ +/+ Δ/+ Δ/Δ

**ADMA FUS**

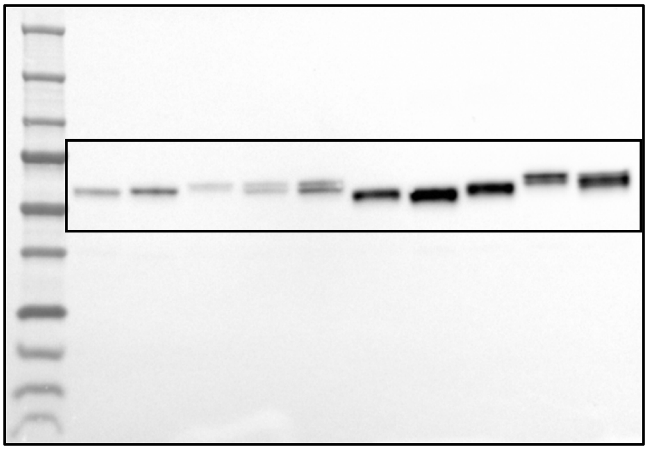

**Cytoplasm** **Nucleus**

No Tg hFUS No Tg hFUS

+/+ Δ/+ +/+ Δ/+ Δ/Δ +/+ Δ/+ +/+ Δ/+ Δ/Δ

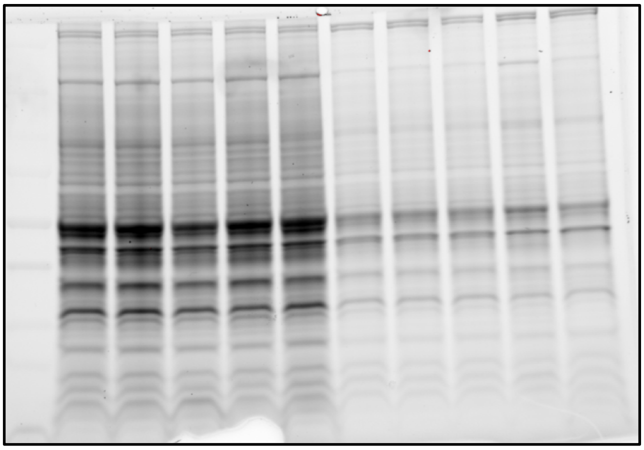

### Sanjuan-Ruiz et al. Figure S4

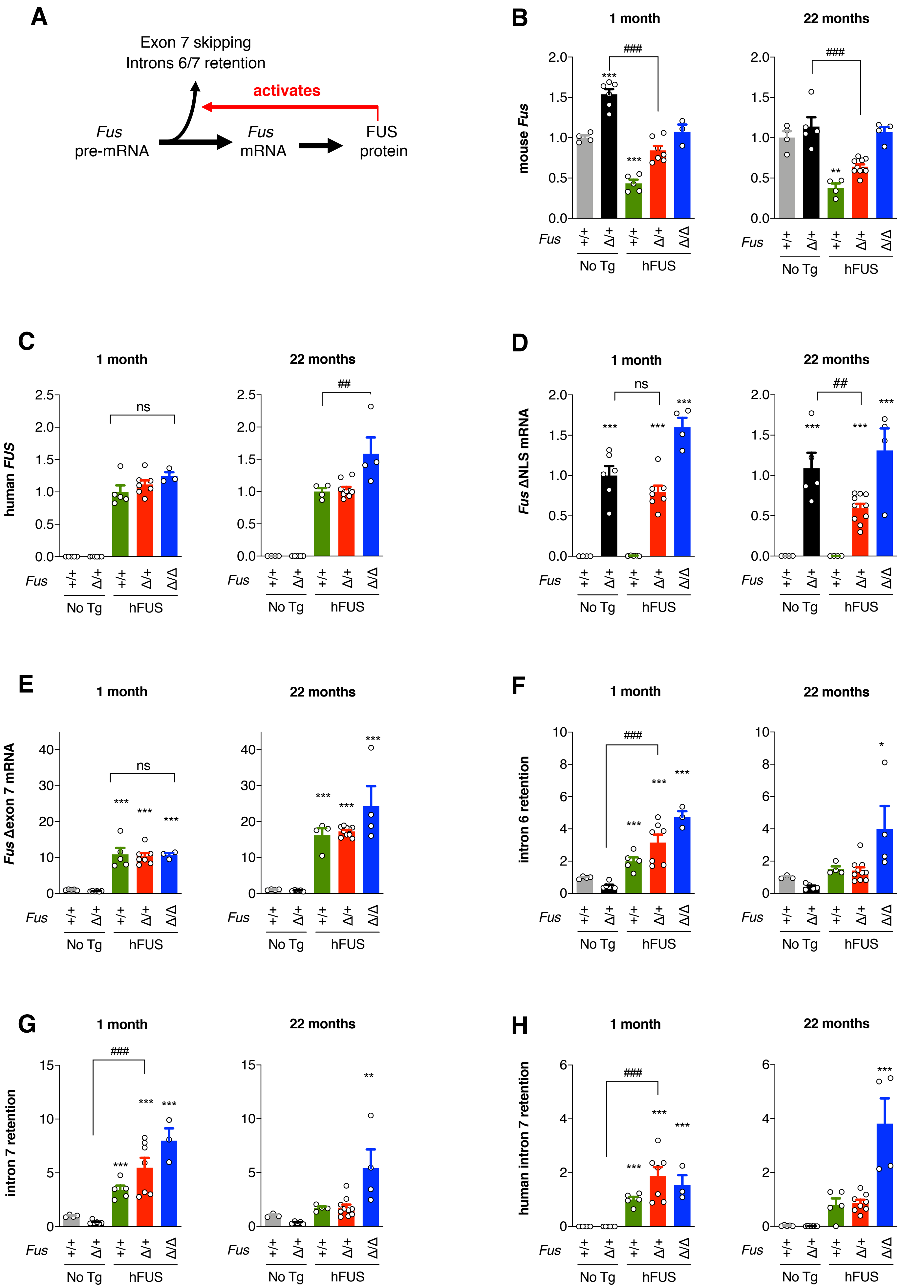

### Sanjuan-Ruiz et al. Figure S5

#### Zeb 1 mRNA levels

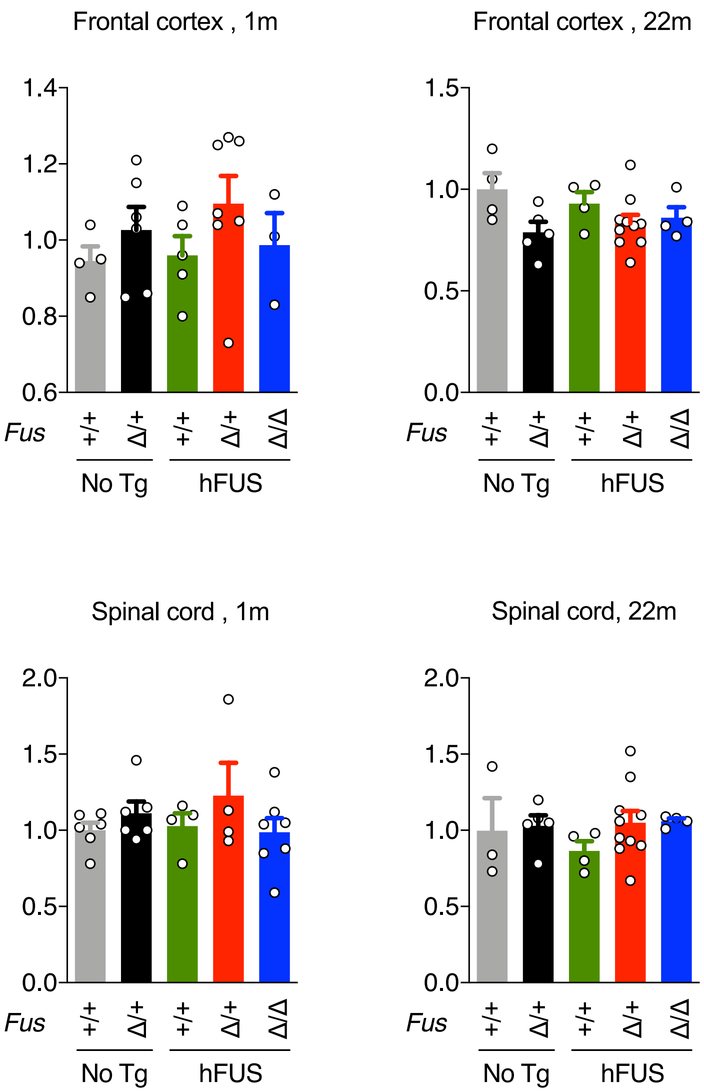
